# Enisamium is a small molecule inhibitor of the influenza A virus and SARS-CoV-2 RNA polymerases

**DOI:** 10.1101/2020.04.21.053017

**Authors:** Alexander P Walker, Haitian Fan, Jeremy R Keown, Victor Margitich, Jonathan M Grimes, Ervin Fodor, Aartjan J W te Velthuis

## Abstract

Influenza A virus and coronavirus strains cause a mild to severe respiratory disease that can result in death. Although vaccines exist against circulating influenza A viruses, such vaccines are ineffective against emerging pandemic influenza A viruses. Currently, no vaccine exists against coronavirus infections, including pandemic SARS-CoV-2, the causative agent of the Coronavirus Disease 2019 (COVID-19). To combat these RNA virus infections, alternative antiviral strategies are needed. A key drug target is the viral RNA polymerase, which is responsible for viral RNA synthesis. In January 2020, the World Health Organisation identified enisamium as a candidate therapeutic against SARS-CoV-2. Enisamium is an isonicotinic acid derivative that is an inhibitor of multiple influenza B and A virus strains in cell culture and clinically approved in 11 countries. Here we show using *in vitro* assays that enisamium and its putative metabolite, VR17-04, inhibit the activity of the influenza virus and the SARS-CoV-2 RNA polymerase. VR17-04 displays similar efficacy against the SARS-CoV-2 RNA polymerase as the nucleotide analogue remdesivir triphosphate. These results suggest that enisamium is a broad-spectrum small molecule inhibitor of RNA virus RNA synthesis, and implicate it as a possible therapeutic option for treating SARS-CoV-2 infection. Unlike remdesivir, enisamium does not require intravenous administration which may be advantageous for the development of COVID-19 treatments outside a hospital setting.

**Importance:** Influenza A virus and SARS-CoV-2 are respiratory viruses capable of causing pandemics, and the latter is responsible for the Coronavirus Disease 2019 (COVID-19) pandemic. Both viruses encode RNA polymerases which transcribe their RNA genomes and are important targets for antiviral drugs including remdesivir. Here, we show that the antiviral drug enisamium inhibits the RNA polymerases of both influenza A virus and SARS-CoV-2. Furthermore, we show that a putative metabolite of enisamium is a more potent inhibitor, inhibiting the SARS-CoV-2 RNA polymerase with similar efficiency to remdesivir. Our data offer insight into the mechanism of action for enisamium, and implicate it as a broad-spectrum antiviral which could be used in the treatment of SARS-CoV-2 infection.

## Introduction

RNA viruses, such as pandemic influenza A viruses (IAV) and severe acute respiratory coronavirus 2 (SARS-CoV-2), are among the most important human pathogens. While IAV and SARS-CoV-2 are different viruses and follow different replication cycles, both can cause severe respiratory disease in humans that leads to high morbidity and mortality. Vaccines exist against influenza viruses; however, long vaccine development times mean that antigenic mismatches can occur between circulating influenza virus strains and the vaccine strain. Moreover, due to antigenic shift, existing vaccines are not effective against emerging pandemic influenza A viruses(1). No vaccine currently exists against coronaviruses, including the SARS-CoV-2 pandemic and SARS-CoV epidemic strains, which cause Coronavirus Disease 2019 (COVID-19) and SARS, respectively. Therefore, research is needed into conserved viral enzymatic activities, such as RNA polymerase activity, which could be targeted by broad spectrum antivirals(2, 3).

IAVs are negative sense RNA viruses whose 14 kb genome consists of eight segments of single-stranded viral RNA (vRNA). The viral RNA-dependent RNA polymerase (FluPol) copies the vRNA into a replicative intermediate called the complementary RNA (cRNA) during viral replication, or into capped and polyadenylated viral messenger RNA (mRNA) during viral transcription(4, 5). The cRNA serves as a template for the production of new vRNA molecules. vRNA and cRNA molecules are both replicated in the context of ribonucleoproteins (RNPs), which consist of FluPol bound to the 5’ and 3’ ends of a genome segment and the rest of the vRNA or cRNA is bound by a helical coil of nucleoprotein (NP). FluPol is composed of three subunits: polymerase basic 1 (PB1), PB2, and polymerase acidic (PA). Structural analyses have shown that the PB1 subunit adopts the canonical polymerase right hand-like fold, which contains the fingers, palm and thumb subdomains that are conserved among all viral RNA polymerases. The PA subunit has a large C-terminal domain that is integrated into the PB1 thumb subdomain, and is connected to an N-terminal endonuclease domain by a linker. The PB2 subunit is composed of several globular domains, including cap binding and 627 domains, which are flexible with respect to the core PB1 subunit(4).

SARS-CoV-2 is a betacoronavirus in the order Nidovirales, and has a positive-sense, non-segmented RNA genome of around 30 kilobases(2, 6). The viral genome has a 5’ m^7^GpppA^m^ cap and 3’ poly(A) tail, modifications which allow the viral genome to be translated by cellular machinery. Two-thirds of the viral genome encodes two overlapping open reading frames (ORFs), 1a and 1b, which are translated into two large polyproteins immediately upon infection. The two polyproteins are cleaved by intrinsic proteolytic activity to produce non-structural proteins (nsps) 1-16, which collectively form the membrane-associated replication-transcription complex (RTC). The RTC has several major functions: Firstly, it synthesises full-length negative-sense RNA antigenomes, which are the templates for new positive-sense RNA genomes. Secondly, it synthesises subgenomic negative-sense RNAs which contain the ribosome-accessible ORFs of structural and accessory proteins. Finally, the RTC transcribes subgenomic or full-length negative-sense RNAs into 5’ m^7^GpppA^m^ capped, 3’ polyadenylated viral mRNAs. The multiple functions of the replicase complex requires nsp1-16 to have many catalytic activities, such as cap synthesis, which are not entirely understood(7). Nsp12 is the RNA-dependent RNA polymerase component of the replicase complex, and it requires nsp7 and nsp8 for processivity(8, 9). The structures of nsp7/8/12 complexes from SARS-CoV and SARS-CoV-2 have been solved by cryo-EM(10–13). Nsp12 is the core of the complex that adopts the canonical right-handed RNA polymerase fold that is linked to an N-terminal domain implicated in nucleotidyltransferase activity(14). One nsp7 and two nsp8 molecules form a complex with each nsp12.

IAV FluPol and SARS-CoV RTC are important drug targets. Favipiravir (T-705) triphosphate (Fig. 1) inhibits FluPol RNA synthesis activity, and other catalytic activities such as the PA endonuclease have also been targeted(15, 16). Remdesivir is a nucleoside analogue drug that has shown promise in both cell culture and clinical trials as a treatment for SARS-CoV-2 infection(17). Remdesivir triphosphate (Fig. 1) is the active metabolite of remdesivir, which inhibits the SARS-CoV-2 nsp7/8/12 complex and is not excised by the nsp14 exonuclease (12). Further nucleoside analogue drugs have also been suggested as therapeutic candidates(18).

**FIG 1.**
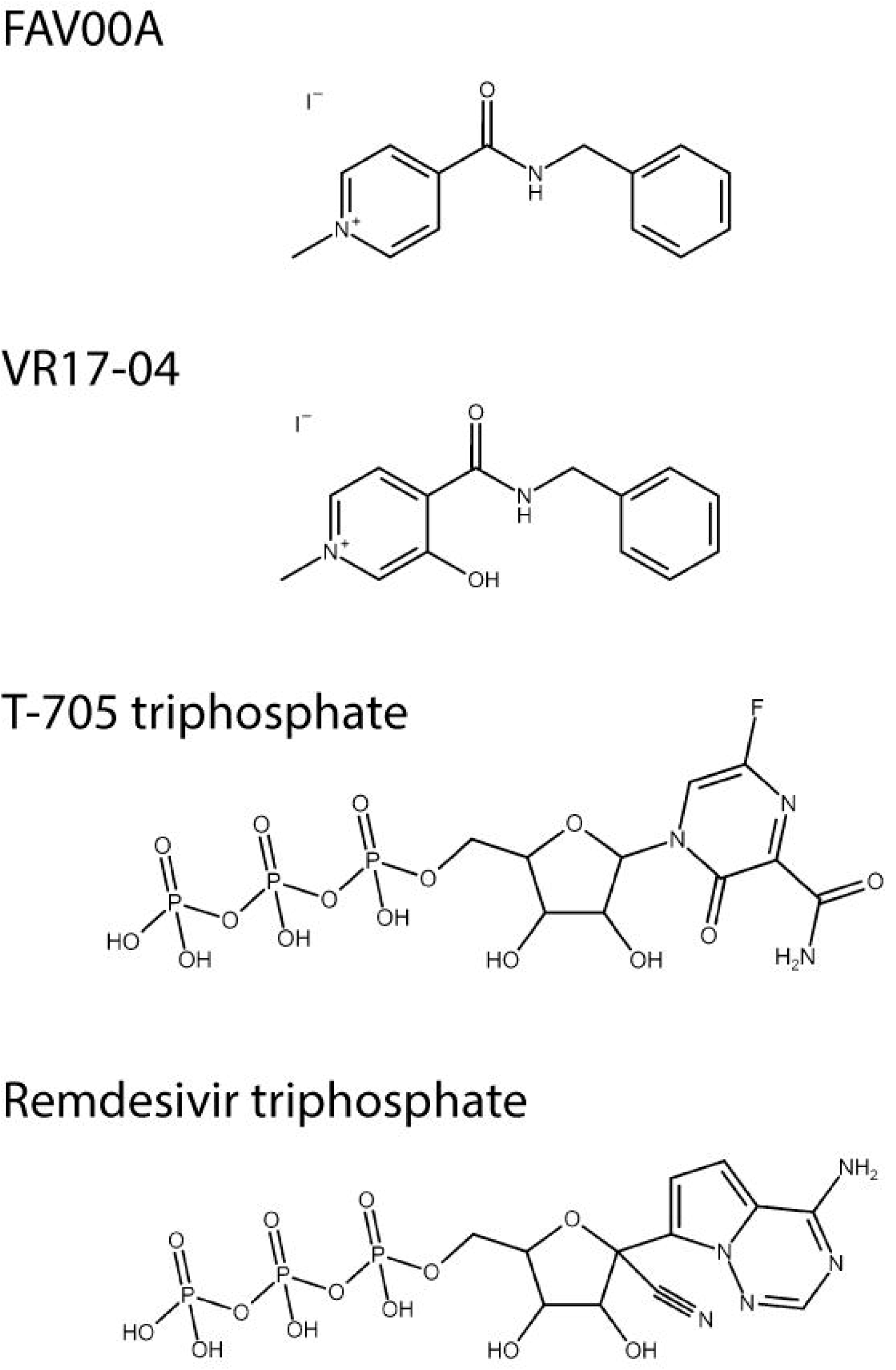
Chemical structures of FAV00A, VM17-04, T-705 triphosphate and remdesivir triphosphate. The structure of FAV00B is identical to FAV00A except that chloride ions are present instead of iodide.

However, there are currently no drugs licensed to treat SARS-CoV-2 infection. Enisamium (4-(benzylcarbamoyl)-1-methylpyridinium) iodide (FAV00A; Fig. 1), marketed as Amizon^®^, was highlighted by the World Health Organisation as a candidate therapeutic against SARS-CoV-2 (19). FAV00A is licensed for use in 11 countries, including Ukraine and Russia, and has been shown to have clinical efficacy against influenza A and B virus infections(20). It is thought to inhibit influenza virus RNA synthesis, however, the mechanism of action is currently unknown(21).

In this study, we show that FAV00A and a putative hydroxylated metabolite (Fig. 1), VR17-04, inhibit FluPol and SARS-CoV-2 polymerase activity *in vitro*. The inhibition of RNA synthesis by both polymerases implies that FAV00A is a broad-spectrum small molecule inhibitor of RNA virus RNA synthesis, and is a possible therapeutic option for treating SARS-CoV-2 infection.

## Results

### Enisamium inhibits FluPol activity in cell culture

Previous experiments showed that enisamium can inhibit influenza virus replication in tissue culture(21). We confirmed these observations by infecting A549 cells with influenza A/WSN/33 (H1N1) virus (WSN) at MOI 0.01 and adding enisamium chloride (FAV00B) to the cell culture medium. Forty-eight hours after infection, virus titres were analysed by plaque assay on MDCK cells. FAV00B significantly affected IAV tires in line with previous observations (Fig. 2A). To test if FAV00B can inhibit IAV RNA synthesis in a minigenome context, HEK 293T cells were transfected with plasmids expressing the PB1, PB2 and PA FluPol subunits, NP and a segment 5 (NP) vRNA template. To estimate the effect of FAV00B on host cell RNA and protein synthesis, we transfected a plasmid expressing GFP from a constitutively active CMV promoter. As a further control, we analysed the effect of enisamium on the 5S ribosomal RNA (rRNA) and tubulin protein steady state level. After transfection, FAV00B was added to the cell culture medium at the concentrations indicated in Fig. 2B. Total cellular RNA was extracted 24 hours post-transfection, and IAV replication (vRNA synthesis) activity was analysed by radioactive primer extension. FAV00B significantly affected IAV replication with an IC_50_ of 354 μM, while there was no significant effect on 5S rRNA, GFP, PA subunit or tubulin expression levels (Fig. 2B). Overall, these observations suggest that enisamium has no effect on host cell viability, but inhibits IAV replication through the inhibition of vRNA synthesis by FluPol.

**FIG 2.**
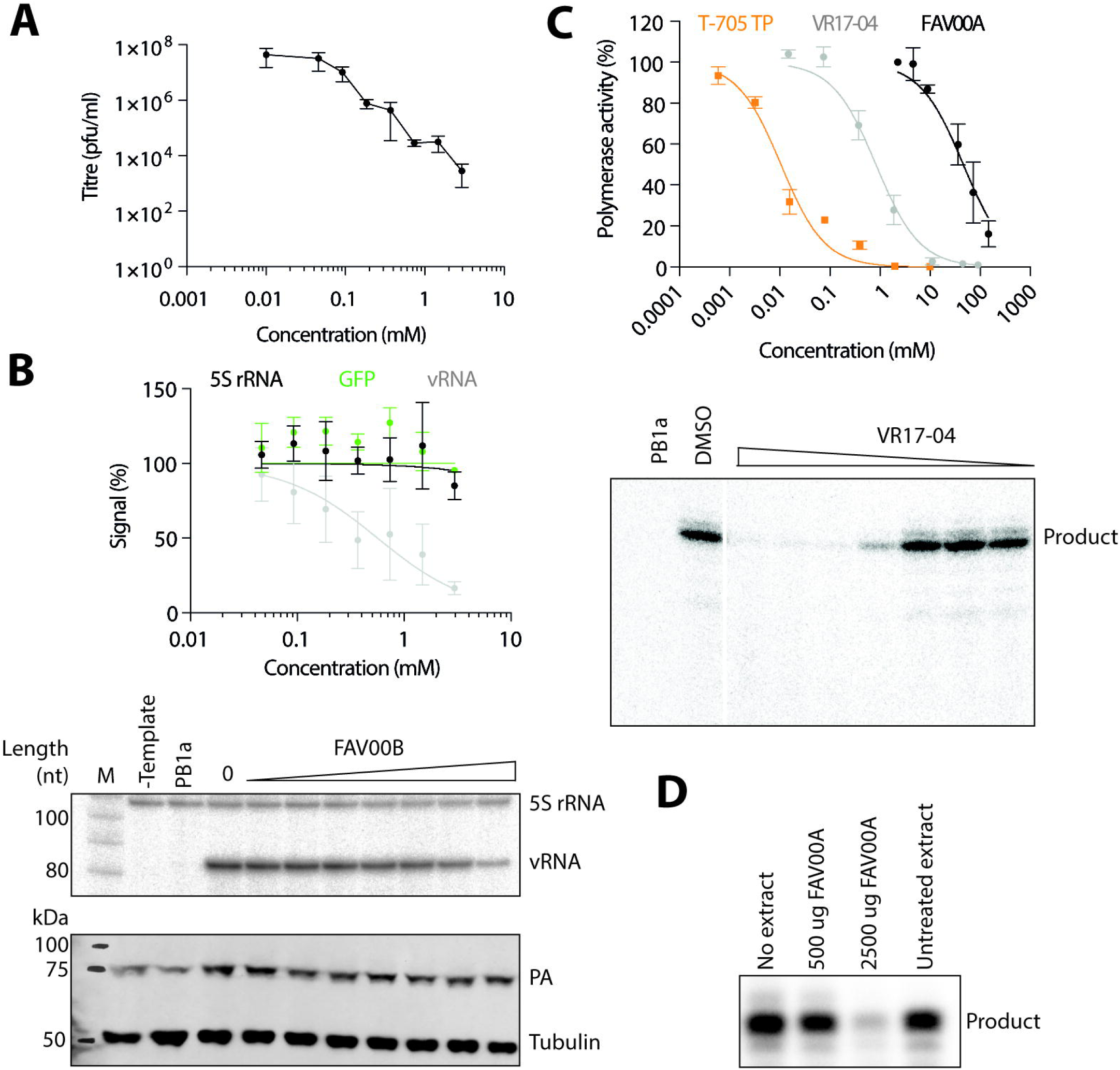
Inhibition of IAV replication and FluPol activity by enisamium. (A) Effect of FAV00B on the replication of influenza A/WSN/33 (H1N1) virus. Data points represent mean and standard deviation of n=3 independent enisamium titrations and matching virus plaque experiments. (B) Effect of FAV00B on the steady-state levels of IAV vRNA, 5S rRNA and the expression of GFP. Levels of 5S rRNA and IAV vRNA were analysed by primer extension (middle panel). PA and tubulin expression were analysed by western blot (bottom panel). A mutant IAV FluPol containing two amino acid substitutions in the PB1 active site (PB1a) was used as negative control. Data points represent mean and standard deviation of n=3 independent enisamium titrations and matching GFP measurements or primer extensions. (C) Effect of T-705 triphosphate, FAV00A and VR17-04 on FluPol activity *in vitro*. Transcription products synthesised by FluPol in the presence of VR17-04 were analysed by PAGE (lower panel). Data points represent mean and standard deviation of n=3 independent enisamium titrations in RNA polymerase assays. (D) Effect of extracts from A549 cells treated with FAV00A on FluPol activity *in vitro*. RNA polymerase products were analysed by PAGE.

### Enisamium and the putative metabolite VR17-04 inhibit FluPol activity

To examine the effect of enisamium on IAV RNA synthesis *in vitro*, we expressed protein A-tagged WSN IAV FluPol in HEK 293T cells and purified the heterotrimeric complex using IgG sepharose chromatography. We then performed *in vitro* RNA synthesis assays using a model 14 nucleotide (nt) viral RNA (vRNA) template in the presence of different concentrations of FAV00A, or T-705 triphosphate as a positive control. In these assays, FAV00A weakly inhibited FluPol activity with an IC_50_ of 46.3 mM, while T-705 triphosphate inhibited FluPol activity with an IC_50_ of 0.011 mM (Fig. 2C). This suggests that the inhibitory effect of enisamium on FluPol activity *in vitro* is limited.

The weak inhibition of FluPol by enisamium is unlikely to be responsible for antiviral efficacy observed in cell culture. To test if a metabolite of enisamium could be a more potent inhibitor of FluPol activity, we treated A549 cells with FAV00A for 24 hours and then performed FluPol activity assays in the presence of cellular extract. We observed a strong, concentration-dependent inhibition of FluPol activity, which is consistent with a metabolite of enisamium inhibiting influenza virus RNA synthesis (Fig. 2D). To test if enisamium metabolites have a direct effect on FluPol activity *in vitro*, 9 metabolites were synthesised (Table 1) and assayed in *in vitro* reactions containing purified FluPol. Only a hydroxylated metabolite, VR17-04, showed substantially increased potency compared to FAV00A (Fig. 1); with an IC_50_ of 0.84 mM it inhibited FluPol activity 55-fold more potently than FAV00A (Fig. 2C).

**Table 1.**
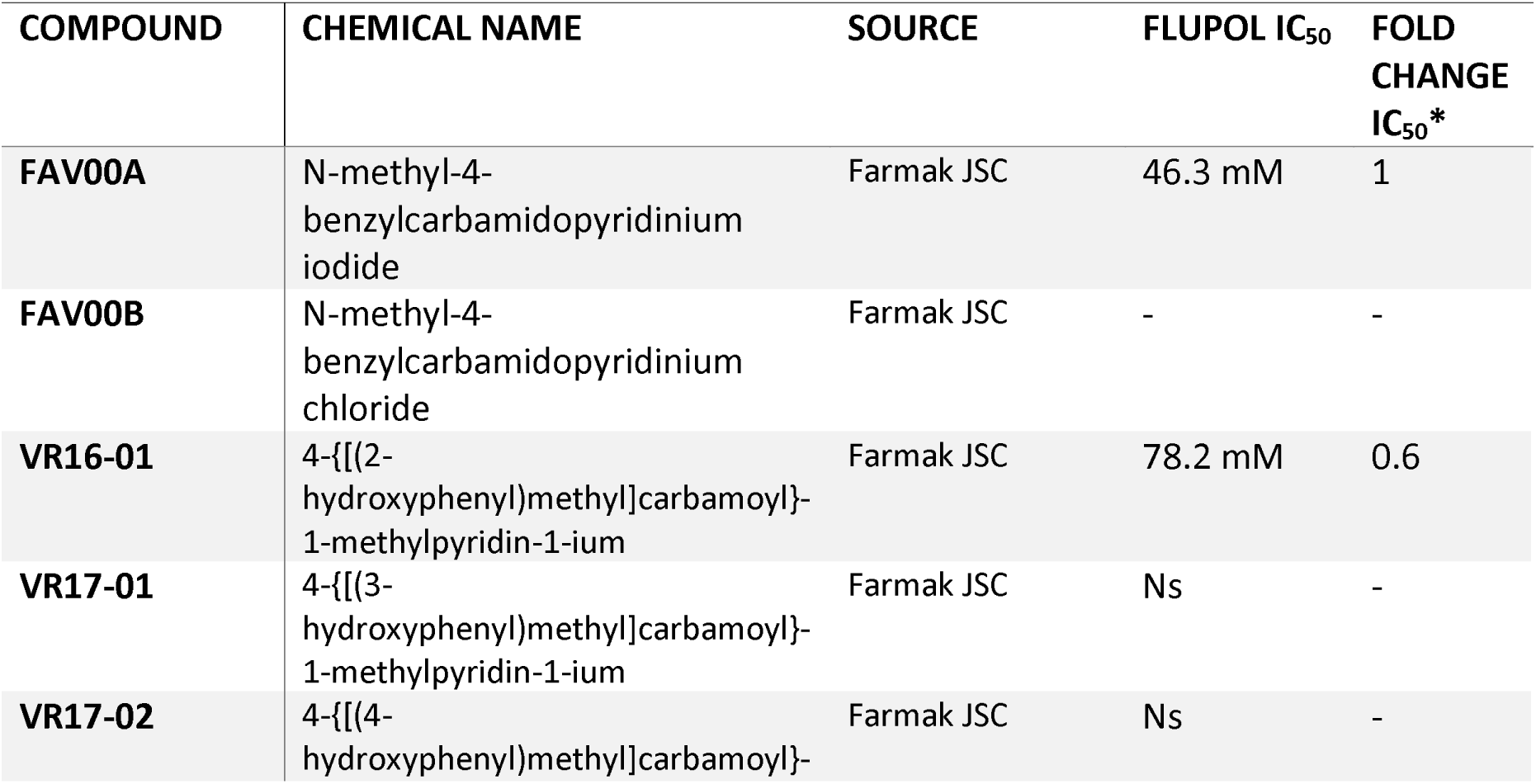
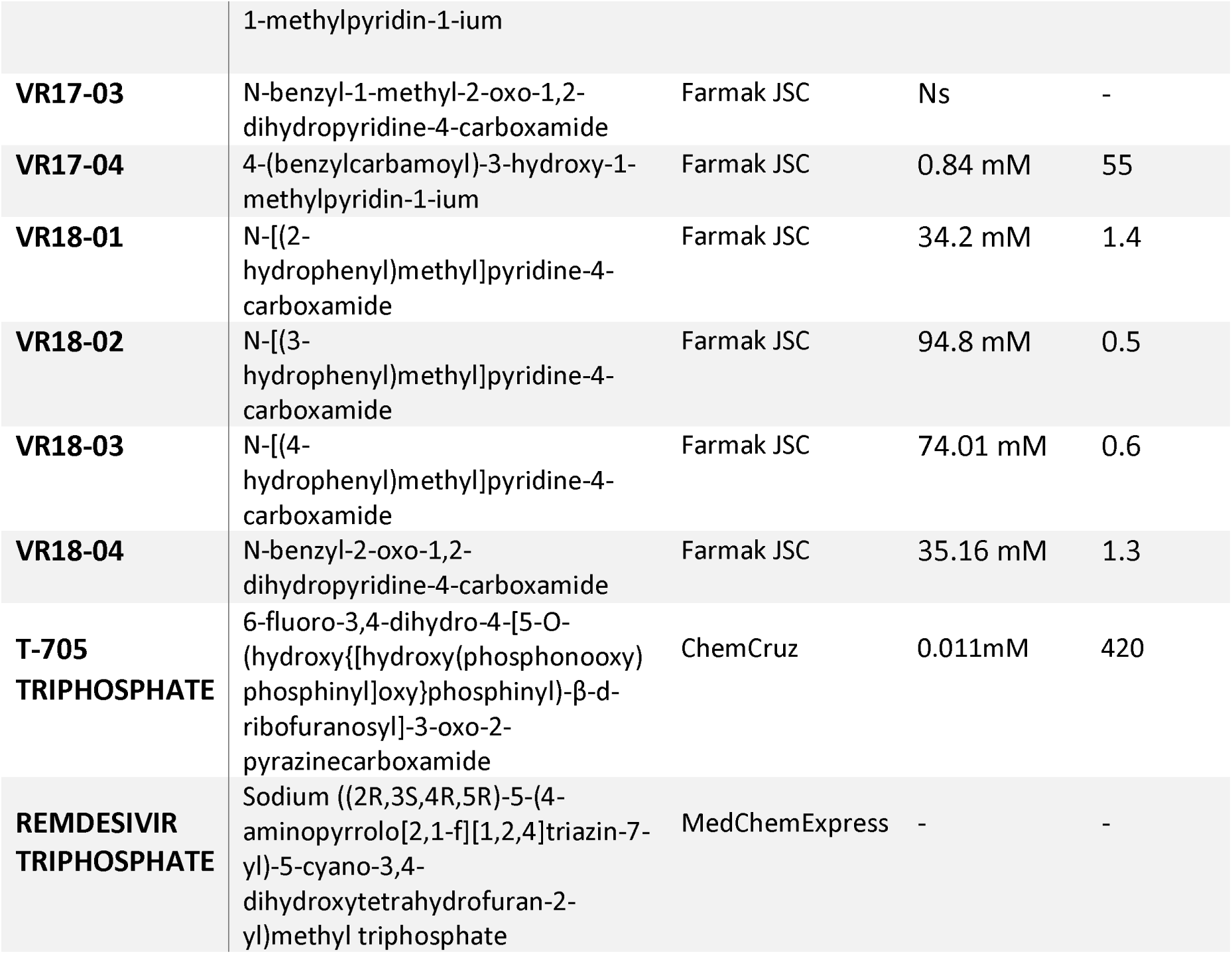
List of compounds tested and their IC_50_ values in FluPol *in vitro* RNA synthesis assays. *Fold change relative to FAV00A. Ns = non-soluble. - = not tested.

To study whether VR17-04 has a direct effect of FluPol initiation or nucleotide incorporation, we measured the effect of VR17-04 on the ability of FluPol to extend a capped RNA primer. In this assay, a capped RNA primer ending in AG can be aligned to the terminal U or UC of the template (Fig. 3A). These two different initiation modes lead to the formation of two extension products, called the C- and G-product (Fig. 3A). A realignment step that can occur as the polymerase slips at 4U of the template can add an additional 3 nt to each extension product, increasing the number of possible extension products(22). Addition of VR17-04 to the capped primer extension assay resulted in absence of extension at high concentrations, and predominantly partial extension of the primer by only 1-2 nucleotides at intermediate VR17-04 concentrations (Fig. 3B). In addition, we observed that formation of the C-product was reduced, suggesting that VR17-04 also affected primer-template binding. Collectively, these data show that hydroxylation of enisamium increases the potency as inhibitor *in vitro* and that hydroxylated enisamium directly affects nucleotide incorporation by FluPol.

**FIG 3.**
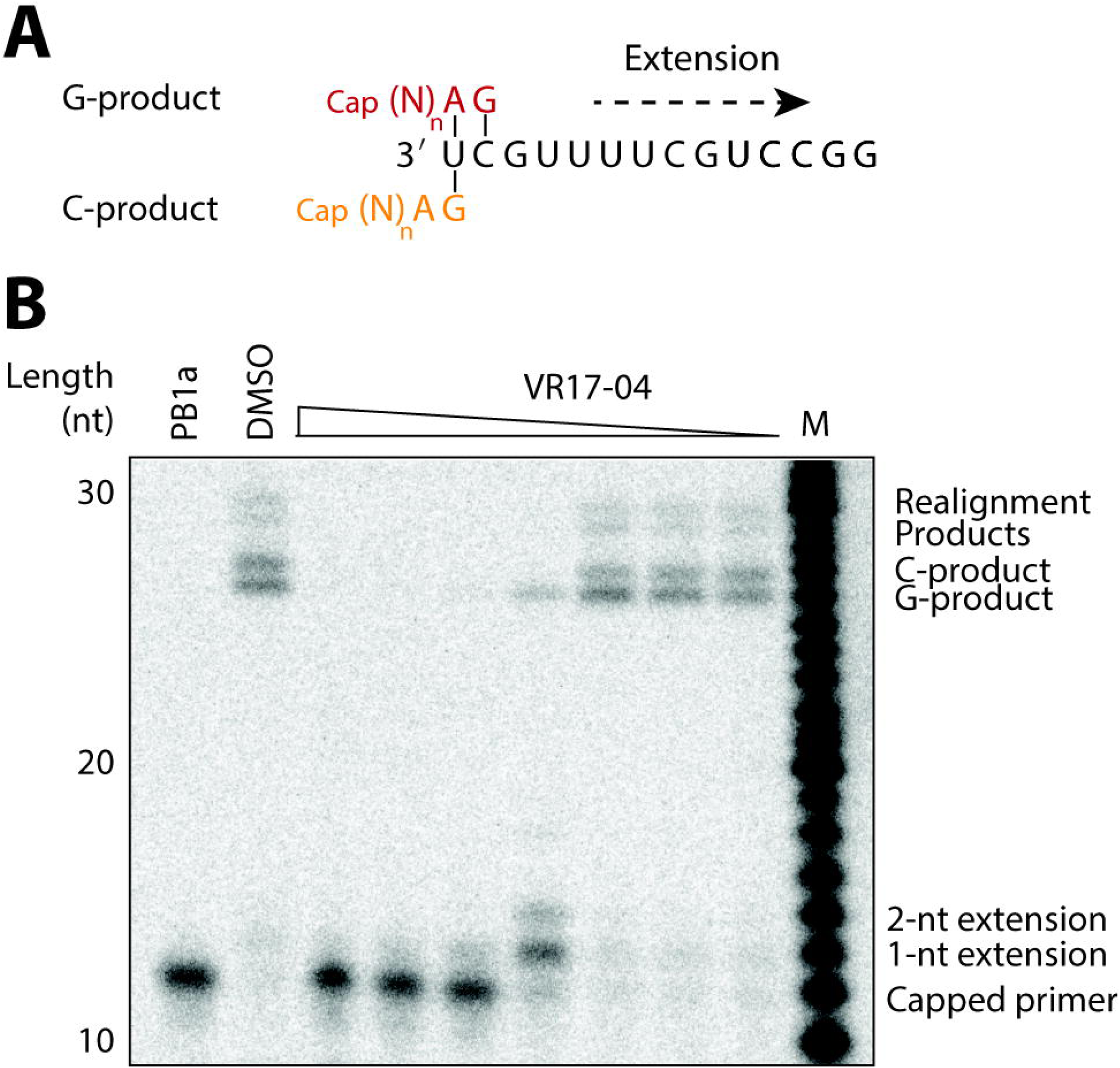
Inhibition of FluPol transcription by VR17-04. (A) Schematic of FluPol transcription initiation using a capped RNA primer. The terminal AG bases of the primer can base pair with the terminal U (orange), or the terminal UC (red) of the influenza virus template, resulting in two possible extension products called the C- and G-product, respectively. (B) Effect of VR17-04 on FluPol transcription *in vitro* using a radiolabelled capped RNA as primer. An active site mutant containing a mutation in the PB1 active site was used as negative control. DMSO was used a solvent control. C- and G-products are indicated as well as realignment products that can form when the polymerase slips at position 4 of the template.

### Enisamium and VM17-04 inhibit SARS-CoV-2 nsp7/8/12 activity

To determine if enisamium and its putative metabolite VR17-04 can inhibit the SARS-CoV-2 RTC, we developed a SARS-CoV-2 RNA polymerase *in vitro* assay that involved nsp12 as the RNA-dependent RNA polymerase, and nsp7 and nsp8 as processivity factors(8). We expressed and purified SARS-CoV-2 nsp7, nsp8 and nsp12, and mixed them at a ratio of 2:2:1 to form a nsp7/8/12 complex (Fig 4A). To test *in vitro* RNA synthesis activity, we incubated the nsp7/8/12 complex with a 40 nt RNA template and a radiolabelled 20 nt RNA primer (Fig 4B, C). The nsp7/8/12 complex was able to extend the 20 nt primer in the presence of NTPs into a doublet of major products of approximately 40 nt in length. This pattern was previously observed for the SARS-CoV nsp7/8/12 complex(8). The proportion of 20 nt primer extended by nsp7/8/12 increased over time, plateauing at around 30 minutes (Fig 4C).

**FIG 4.**
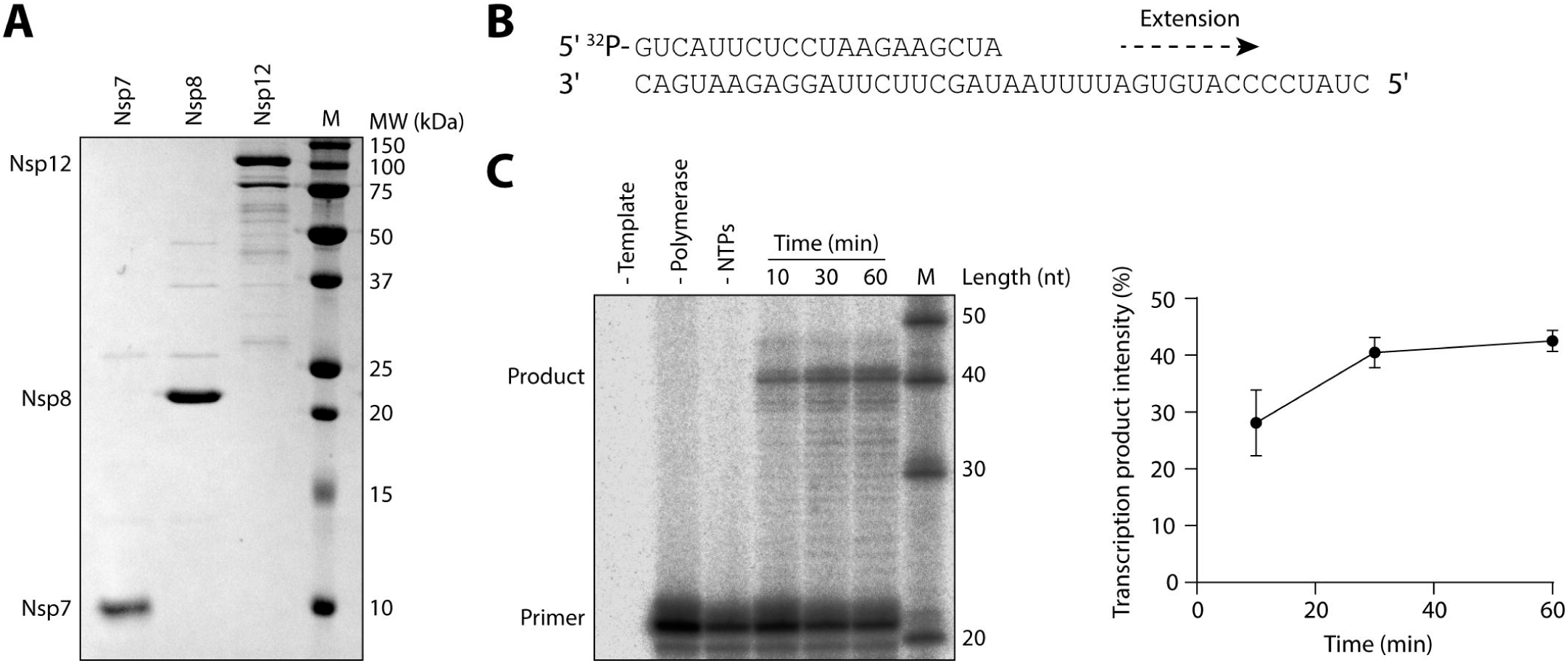
Purification and RNA synthesis activity of SARS-CoV-2 nsp7/8/12 complex. (A) SARS-CoV-2 nsp7, nsp8 and nsp12 were expressed and purified from E. coli (nsp7 and nsp8) or Sf9 (nsp12) cells and visualised by SDS PAGE. (B) Schematic of 40 nt RNA template and 20 nt radiolabelled primer. (C) Purified nsp7/8/12 complex was tested for *in vitro* activity in the presence or absence of NTPs and RNA template. The 20 nt primer and 40 nt product ran slightly slower than the markers, possibly due to differences in the RNA sequence or phosphorylation state. Quantification of the transcription products is shown on the right. Quantification is from n=3 independently prepared reactions using the same nsp7/8/12 protein preparation, error bars represent standard deviation.

Remdesivir triphosphate is a nucleotide analogue shown to inhibit SARS-CoV-2 nsp7/8/12 activity(12). To estimate the potency of this inhibition, we performed *in vitro* RNA synthesis assays in the presence of remdesivir triphosphate or T-705 triphosphate (Fig 5A). Remdesivir triphosphate inhibited RNA synthesis activity with an IC_50_ of 2.01mM, while T-705 triphosphate was non-inhibitory at all concentrations tested. Next, we performed *in vitro* RNA synthesis assays in the presence of FAV00A and the putative metabolite VR17-04 (Fig 5B). FAV00A inhibited nsp7/8/12 activity at relatively high concentrations, with an IC_50_ of 40.7 mM. VR17-04 exhibited over 10-fold more potent inhibition, with an estimated IC_50_ of 2-3 mM, similar to that obtained for remdesivir triphosphate. These IC_50_ values are within an order of magnitude of those calculated for the inhibition of FluPol by FAV00A and VR17-04 (Fig 5C). Collectively, these data show that RNA synthesis by the SARS-CoV-2 nsp7/8/12 complex is inhibited by enisamium and its putative metabolite VR17-04.

**FIG 5.**
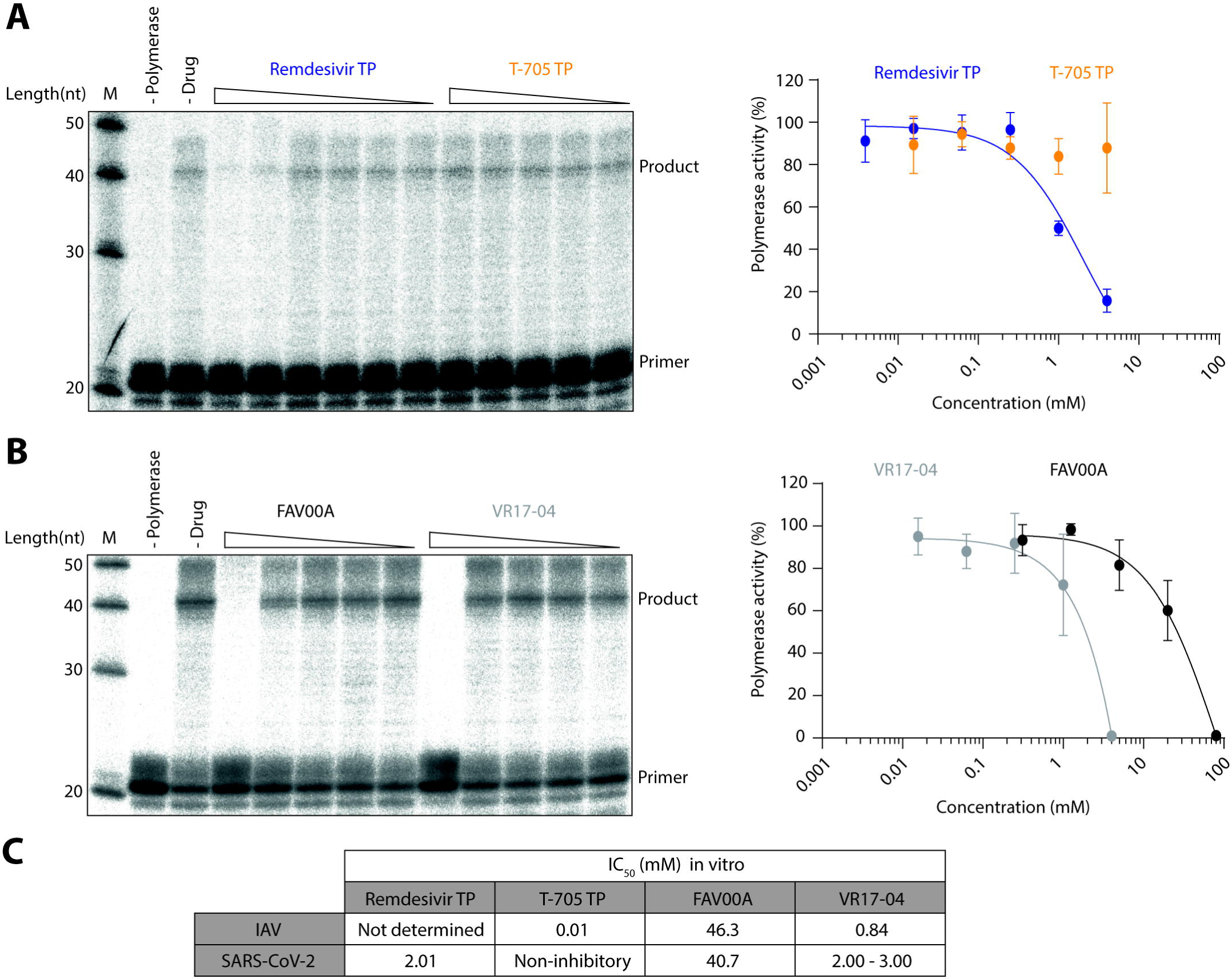
Inhibition of SARS-CoV-2 nsp7/8/12 by remdesivir triphosphate, T-705 triphosphate, enisamium and VR17-04. (A) Effect of remdesivir triphosphate and T-705 triphosphate on nsp7/8/12 activity *in vitro*. Polymerase activity was plotted against drug concentration and dose-response curves were fit to the data (right). (B) Effect of FAV00A and VR17-04 on nsp7/8/12 activity *in vitro*. Polymerase activity was plotted against drug concentration and dose-response curves were fit to the data (right). Quantification is from n=3-6 independently prepared reactions using the same nsp7/8/12 protein preparation, error bars represent standard deviation. (C) IC_50_ values of remdesivir triphosphate, T-705 triphosphate, FAV00A and VR17-04 for IAV FluPol and SARS-CoV-2 nsp7/8/12. The IC_50_ value for VR17-04 on SARS-CoV-2 nsp7/8/12 could not be calculated and therefore it was estimated from the dose-response curve.

## Discussion

In this study we showed that enisamium and a putative metabolite, VR17-04, inhibit FluPol RNA synthesis activity *in vitro*. We also established an *in vitro* RNA synthesis assay for the nsp7/8/12 complex of SARS-CoV-2, and used this assay to show that enisamium and VR17-04 also inhibit RNA synthesis by SARS-CoV-2 nsp7/8/12. Specifically, we find that VR17-04 inhibits SARS-CoV-2 RNA synthesis with a similar IC_50_ to remdesivir triphosphate.

Our *in vitro* assays indicate that FAV00A inhibits FluPol activity with a relatively high IC_50_ value of 46.3 mM (Fig 2C). This inhibition was improved 55-fold by addition of a hydroxyl group in the compound VR17-04 (Fig. 1), which we propose is an active metabolite of enisamium. Similar IC_50_ values of 40.7mM and 2-3mM were observed for FAV00A and VR17-04, respectively, against the SARS-CoV-2 RNA polymerase (Fig 5C). The exact mechanism of RNA synthesis inhibition by enisamium is unknown, but our *in vitro* assays suggest that enisamium can directly inhibit FluPol elongation (Fig. 3). Further studies, such as drug binding assays, could be used to provide further insight into the mechanism of action of VR17-04.

While VR17-04 inhibits FluPol activity with an IC_50_ of 0.84 mM, enisamium inhibits influenza virus growth in cell culture more potently with EC_90_ values ranging from 157-439 µM (Fig. 2A)(21). This apparent discrepancy may be due to enisamium exerting additional antiviral effects through other mechanisms or more potent unknown metabolites. However, it could also be due to the *in vitro* assays not fully reflecting the requirements for RNA polymerase activity in an *in vivo* situation. While our FluPol *in vitro* assays examine the synthesis of a short 14 nt product, in vivo, FluPol must processively synthesise RNA products ranging in length between 890 nt and 2341 nt depending on the viral genome segment(23). In the SARS-CoV-2 nsp7/8/12 *in vitro* activity assay we examine the extension of an RNA primer by 20 nt, however, during infection, nsp7/8/12 must synthesise RNA products up to 30 kilobases(7). Moreover, protein concentrations in the *in vitro* reactions may be higher than in cell culture systems. Therefore, it is not surprising that RNA synthesis inhibitors can show better efficacy *in vivo* than *in vitro*. Indeed, this phenomenon is exemplified by the nucleoside analogue drug remdesivir. Remdesivir has a reported IC_50_ of 0.77 µM against SARS-CoV-2 in cell culture, while we find that remdesivir triphosphate has a much higher IC_50_ of 2.01 mM *in vitro*, which is similar to values observed previously (Fig 5A)(12, 17).

We find that SARS-CoV-2 nsp7/8/12 complex activity is inhibited by FAV00A and VR17-04 with similar IC_50_ values to FluPol, 40.7mM and 2-3mM respectively (Fig 5C). This suggests that the concentration of enisamium required for *in vivo* efficacy against SARS-CoV-2 may be similar to that required for influenza virus, which is not toxic in cell culture and humans (21)(Fig. 2B). Furthermore, the IC_50_ value of VR17-04 against nsp7/8/12 is similar to that of remdesivir triphosphate (Fig 5C). The similar *in vitro* efficacy of remdesivir triphosphate and the putative enisamium metabolite VR17-04 suggests that enisamium could inhibit SARS-CoV-2 replication as well.

The rapid global spread of SARS-CoV-2 necessitates development of effective therapeutic interventions, and the most promising short-term strategy is to repurpose existing drugs. Our results demonstrate that enisamium, which is approved for use against influenza in 11 countries, inhibits RNA synthesis by the SARS-CoV-2 nsp7/8/12 complex. A putative enisamium metabolite has similar *in vitro* efficacy against SARS-CoV-2 RNA polymerase activity as remdesivir triphosphate. This raises the possibility that enisamium could be a viable therapeutic option against SARS-CoV-2 infection. Moreover, unlike remdesivir, enisamium does not require intravenous administration, which would be advantageous for its use outside of a hospital setting. A randomised, double-blind, placebo-controlled, parallel-group phase 3 trial will need to be used to assess the antiviral efficacy and safety of enisamium iodide (Amizon^®^) in subjects with moderate COVID-19. Indeed, permission is currently being sought for a phase 3 clinical trial of Amizon^®^ Max for the treatment of COVID-19 in Ukraine.

## Materials and Methods

### IAV infections

A549 cells were preincubated with varying concentrations of FAV00B in Minimal Essential Medium (MEM) containing 0.5% foetal calf serum (FCS) at 37 °C for 1 h. Cells were next infected with influenza A/WSN/33 (H1N1) virus at a multiplicity of infection (MOI) of 0.01 in MEM containing 0.5% FCS for 1 h. Following infection, the inoculum was removed and replaced with varying concentrations of FAV00B in MEM containing 0.5% FCS and incubated for 48 h. Plaque assays were performed on MDCK cells in MEM containing 0.5% FCS with a 1% agarose overlay and grown for 2 days at 37 °C.

### Minigenome assays

HEK 293T cells grown in Dulbecco’s Minimal Essential Medium (DMEM) with 10% FCS were transfected with plasmids expressing influenza A/WSN/33 (H1N1) virus RNA polymerase subunits (pcDNA3-PB1, pcDNA3-PB2, pcDNA3-PA), NP (pcDNA3-NP) and segment 5 (NP) vRNA (pPOLI-NP-RT)(24, 25). As negative control, a PB1 active site mutant (PB1a) was used in which the active site SDD motif was mutated to SAA(26). GFP was expressed through transfection of pcDNA3-EGFP(27). FAV00B was added to the cell culture medium 15 min after transfection. Twenty-four hours after transfection, cells were washed in PBS and the total RNA extracted using Tri-Reagent (Sigma) and isopropanol precipitation. IAV vRNA and 5S rRNA steady state levels were subsequently analysed by radioactive primer extension and 6% denaturing PAGE as described previously(28). Phosphorimaging was performed on a Typhoon Scanner. For GFP measurements, approximately 0.2×10^6^ cells were suspended in 100 μl PBS and transferred to a black 96-well Cellstar plate (Greiner Bio-One). Fluorescence was measured on a SpectraMax i3 plate reader using standard settings. Western analysis was performed on 8% SDS-PAGE and PA antibody GTX125932 (GeneTex)

### FluPol *in vitro* activity assays

The subunits of the A/WSN/33 (H1N1) influenza virus FluPol with a Tandem Affinity Purification (TAP)-tag at the C-terminus of PB2 (pcDNA3-PB1, pcDNA3-PB2-TAP, pcDNA3-PA) were expressed in HEK 293T cells and purified as heterotrimeric complex using IgG sepharose chromatography as described previously(29). Next, 0.5 mM CTP, 0.5 mM UTP, 0.05 mM ATP, 0.01 mM GTP, 0.5 mM ApG, 0.001 mM [α-^32^P]GTP, T-705 triphosphate, FAV00A and VR17-04 at the concentrations indicated, and 0.5 μM 5’ (5’-AGUAGUAACAAGGCC-3’)and 3’ (5’-GGCCUGUUUUUACU-3’) vRNA promoter strands were added to purified FluPol. After incubation at 30 °C for 15 min, samples were analysed by 20% denaturing PAGE and autoradiography. Transcription assays were performed as described previously, using radiolabelled capped 11-nucleotide RNA (5⍰-pppGAAUACUCAAG-3⍰; Chemgenes)(22, 30).

### FAV00A-treated A549 cell extracts

Approximately 10^6^ A549 cells treated with FAV00A in DMEM containing 10% FCS were lysed through sonication in 40 μl buffer A (10 mM HEPES-NaOH, pH 7.5, 0.1% NP-40, 150 mM NaCl, 5% glycerol, and 1 mM DTT) for 10 seconds. After centrifugation, extract was added to purified FluPol reactions at 1/10 of the total reaction volume.

### SARS-CoV-2 protein expression and purification

Full-length nsp7, nsp8, and nsp12 genes from SARS-CoV-2, codon optimized for insect cells, were purchased from IDT. The nsp12 gene was cloned into the MultiBac system, with a Tobacco Etch Virus (TEV) protease cleavable protein-A tag on the C-terminus(31). The nsp7 and nsp8 genes were cloned into pGEX-6P-1 vector (GE Healthcare) with an N-terminal GST tag followed by a PreScission protease site. Nsp12 was expressed in Sf9 insect cells and nsp7 and nsp8 were expressed in E. coli BL21 (DE3) cells. Initial purification of nsp12 was performed by affinity chromatography as previously described for FluPol with minor modifications: all buffers were supplemented with 0.1 mM MgCl_2_ and the NaCl concentration was changed to 300 mM(32). Nsp7 and nsp8 were purified on Glutathione Sepharose (GE Healthcare). After overnight cleavage with TEV (nsp12) or PreScission (nsp7 and nsp8) proteases, the released proteins were further purified on a Superdex 200 (for nsp12) or a Superdex 75 (for nsp7 and nsp8) Increase 10/300 GL column (GE Healthcare) using 25 mM HEPES–NaOH, pH 7.5, 300 mM NaCl and 0.1 mM MgCl_2_. Fractions of target proteins were pooled, concentrated, and stored at 4 °C.

### SARS-CoV-2 nsp7/8/12 *in vitro* activity assays

Nsp7, nsp8 and nsp12 were mixed at a molar ratio of 2:2:1 and incubated overnight at 4 °C to form the nsp7/8/12 complex. Activity assays were performed essentially as described previously for SARS-CoV-2 nsp7/8/12(8). Briefly, 40mer (5’-CUAUCCCCAUGUGAUUUUAAUAGCUUCUUAGGAGAAUGAC-3’) and radiolabelled 20mer (5’-GUCAUUCUCCUAAGAAGCUA-3’) RNAs corresponding to the 3’ end of the SARS-CoV genome (without the polyA tail) were pre-annealed by heating to 70 °C for 5 mins followed by cooling to room temperature. 50 nM pre-annealed RNA was incubated for 10-60 mins at 30 °C with 500 nM nsp7/8/12 complex, in reaction buffer containing 5 mM MgCl_2_, 0.5 mM of each ATP, UTP, GTP and CTP, 10 mM KCl, 1 U RNasin (Promega) and 1 mM DTT. Assays in the presence of remdesivir triphosphate, T-705 triphosphate, FAV00A or VR17-04 were run for 30 mins. Reactions were stopped by addition of 80 % formamide and 10 mM EDTA, followed by heating to 95 °C for 3 mins. Reaction products were resolved by 20 % denaturing PAGE with 7M urea, and visualised by phosphorimaging on a Fuji FLA-5000 scanner. Data were analysed using ImageJ and Prism 8 (GraphPad).

## Acknowledgments

A.J.W.t.V is supported by joint Wellcome Trust and Royal Society grant 206579/Z/17/Z and the National Institutes of Health grant R21AI147172. E.F. is supported by Medical Research Council program grant MR/R009945/1. J.M.G. is supported by Wellcome Investigator Award 200835/Z/16/Z. A.P.W. is supported by Medical Research Council studentship 1960073. This work was also supported by the COVID-19 Research Response Fund administered through the Medical Sciences Division.

All authors contributed to the design and conceptualisation of the work. A.P.W., A.J.W.t.V., H.F. and J.K. performed the experiments and analyzed the data. V.M. provided compounds. A.P.W. and A.J.W.t.V. wrote the paper with contribution from all authors.

## Conflict of interest

V.M. is an employee of Farmak Public Joint Stock Company, Kiev, Ukraine. Part of this research was funded by Farmak Public Joint Stock Company, Kiev, Ukraine.

## Notes

### Summary of Updates

Additional data added: influenza virus infection in A549 cells; comparisons with favipiravir and remdesivir in vitro; effect vr17-04 on influenza virus transcription in vitro. These data appear in Fig. 2, 3 and 5.

